# Eye movements reveal representational geometry

**DOI:** 10.64898/2026.07.21.739743

**Authors:** Thérèse Collins

**Affiliations:** Université Paris Cité, CNRS, Integrative Neuroscience and Cognition Center, F-75006 Paris, France

## Abstract

Mental representations are the explanatory construct of the cognitive sciences, but there is no widely accepted characterization of how they cause behavior. Visual representational geometry can be quantified by similarity scores, but almost all methods require an explicit judgment. To examine how representations cause behavior by varying task demands, observers must perform different tasks while continuously reporting similarity, leading to dual-task interference. This study develops scanpaths as an implicit similarity measure, and uses representational similarity analysis to validate it. Observers searched for a target; fixations on distractors may reveal similarity. Similarity was also quantified by an odd-one-out task in the same participants, and ratings from different participants (Jiang et al. 2022). Representational geometries between tasks correlated. A generative model predicted first fixations in novel data. This double validation of the scanpath method opens the door to examining the causality of representations by determining if and how they vary with task demands.

## Introduction

Representations are the central explanatory construct of the cognitive sciences because they link internal states to perception, memory, and behavior. Their content is defined by the relationships among representational states – that is, by representational geometry (Shepard, 1962; Shepard & Chipman, 1972; Shepard, 1980; Edelman, 1998; Baker et al., 2022). Yet representational geometry only earns explanatory value if it explains why we act in a particular way (James, 1890). Measuring representations independently of the behaviors they are meant to explain therefore leaves a fundamental methodological and theoretical gap. Thus, while it is crucial to recover representational geometry directly from behavior as it unfolds, most methods do not allow this. Traditionally, cognitive psychology has quantified representations by asking observers to rate how similar objects or concepts are to one another either with straightforward similarity scores, or by odd-one-out or arrangement tasks (Kriegeskorte & Mur, 2012). All of these tasks require participants to perform an explicit similarity task, which makes it difficult to examine the representations that may drive another behavior that probes perception or memory, for obvious dual-task interference problems: it is not possible to ask participants to simultaneously perform a task while also continuously reporting similarity.

Eye movement guidance during visual search and scene viewing has long been described as a competition between objects on a saliency or priority map (Wolfe, 2020; Henderson, 2017). Which objects are fixated by the eyes or attended to next depends on their weight in such a map. Maps can combine both bottom-up salience and top-down target similarity or task-relevance to determine the next most relevant location for attention and/or the eyes to move to. In visual search, distractors that are similar to the target along a number of possible features or dimensions are highly activated and are therefore more likely to attract attention and saccades. In scene viewing, semantically relevant regions of a scene are highly activated and attract the eyes and attention more (Henderson & Hayes, 2017; 2018; Hayes & Henderson, 2021). Thus, eye movements during visual search are behaviorally observable consequences of a kind of similarity computation between internal goals or target templates and candidate scene objects (Becker, 2010; Becker et al. 2020). If indeed visual search is like a graded competition between items in a representational space, the question remains open as to whether this behavior is tapping into the same representations as those traditionally considered in explicit similarity tasks, on which the majority of studies on representational geometry rely on. If gaze allocation functions as an indirect measure of representational proximity, this would open the door to using them while observers are engaged in another task, to determine how representations shape behavior as it unfolds.

There has been a recent resurgence of interest in both mental and neural representational geometry in cognitive neuroscience, but most studies do not include a behavioral task, or only a single task, or use an offline task in different participants (Rademaker, Chunharas & Serences, 2019; Chunharas, Wolff, Hettwer & Rademaker, 2025; Walsh & Rissman, 2023; Hebart, Contier, Teichmann, Rockter, Zheng, Kidder, … & Baker, 2023). These studies explain their given set of items in the context of a single task, or describe the geometry of neural representations, without linking them to ongoing behavior. They thus fall short of discovering the functional dimensions of visual experience because they ignore how representations support dynamic behavior.

The current study set out to determine whether eye movements contain latent information about representational geometry. Observers performed a visual search task while their eye movements were continuously monitored. The probability of fixating an item while searching for a target was taken as a measure of their similarity. The similarity of the same set of items was also measured in a triplet odd-one-out task, performed by the same observers, and by a direct comparison score, obtained from different observers in the dataset gathered and made public by Jiang, Sanders & Cowell (2022). Using representational similarity analysis (Kriegeskorte, Mur & Bandettini, 2008), the correlation between tasks was used to establish convergent validity between measures.

Criterion validity (the extent to which the method is predictive of future behavior) was established by a generative model. Generative models specify the internal processes that give rise to outputs (in contrast to predictive models whose aim is to correctly quantify outputs, without saying why those outputs occur). If psychological representations drive oculomotor behavior, then once the representational geometry of a set of items is known, it should be possible to predict which item will be fixated in new data.

## Methods

### Participants

Forty-four participants (31 females, ages 21±41) were recruited from a French national subject pool maintained by the RISC (Relais d’Information sur les Sciences de la Cognition, UAR 3332 CNRS). They came to the laboratory for two separate sessions, in counterbalanced order (visual search task, triplet task), and were compensated for their time as per laboratory practices. All participants signed a consent form. The experiment was approved by the French national ethics board (Comité de Protection des Personnes).

### Stimuli

All images were color photographs of animals, taken from the public dataset made public by Jiang et al. (2022). The full description of these stimuli can be found in their Methods. Stimuli presentation was controlled with Psychtoolbox for MATLAB (MathWorks; Brainard, 1997; Pelli, 1997; Kleiner et al., 2007).

### Visual search task

Participants were informed that they would have to search for a target that was identified by its name (in French), written at screen center. Word length varied but height was roughly 2 dva (depending on exact letters), i.e. words were highly visible. Each image subtended 1.5 dva on screen. Each trial had 13 animal images, the target plus 12 distractors (Figure 1A). Each animal image was selected as the target three times, and the remaining 12 distractors were selected to optimize the coverage of each target-distractor pair. Their positions were chosen randomly with the constraints that no image could be presented within 5 dva of screen edges, and the edges of two images had to be separated by at least 1 dva.

**Figure 1.**
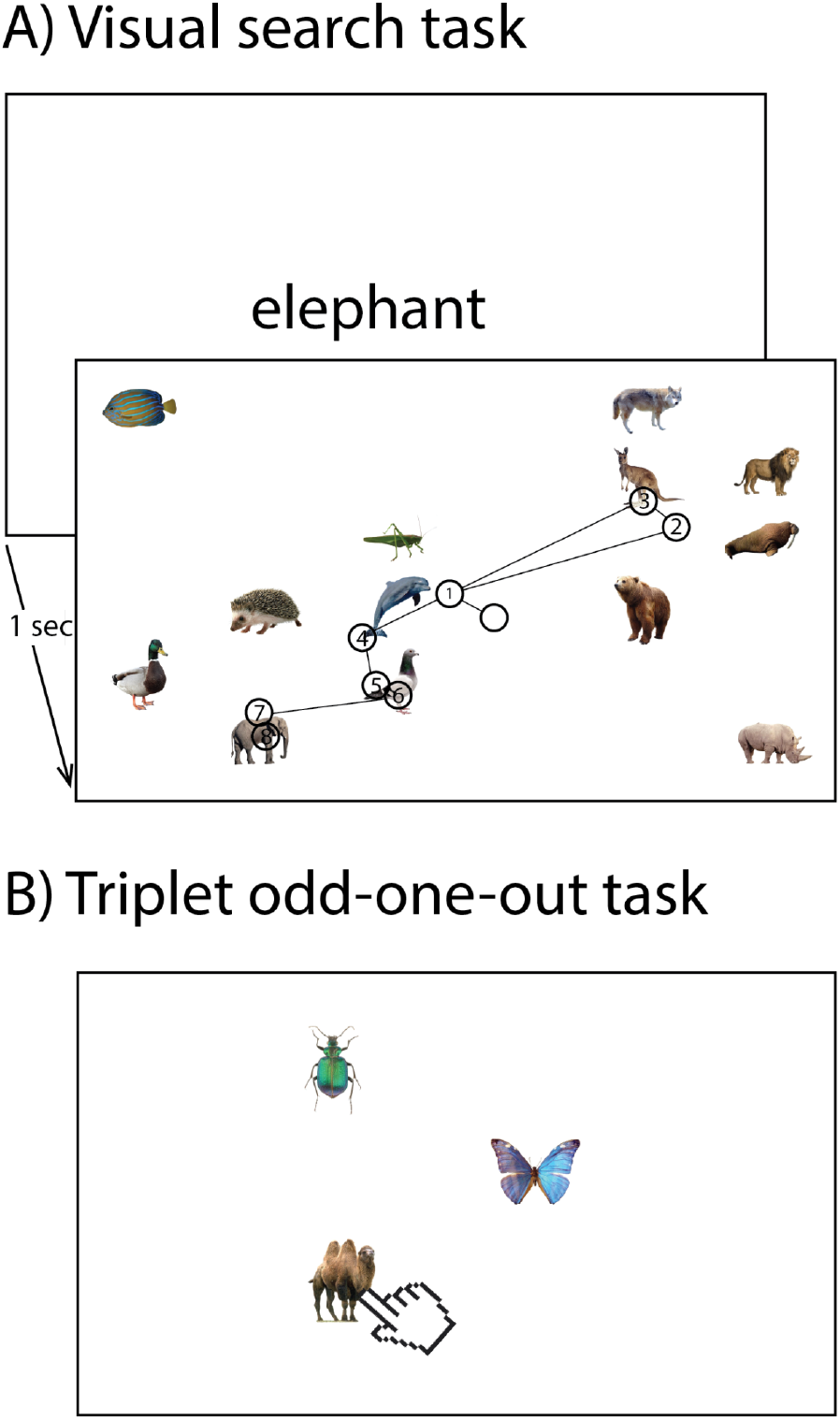
A) Visual search task. The target name was presented for 1 second, followed by the target + distractor screen. Fixations are open circles with a number indicating their rank. Once the target had been found, observers were instructed to maintain fixation on it and press the space bar. Gaze always started at screen center. The example illustrated here shows eight successive fixations that ended up on the target at the bottom right. B) Triplet odd-one-out task. An example triplet trial with beetle, butterfly, camel, in which the observer selected the camel as most different from the two others.

### Triplet odd-one-out task

On each trial, participants were presented with three animal images, each subtending 4 dva, placed on an imaginary 5 dva-radius circle at 0, 135 and 270 degrees (Figure 1B).

### Direct comparison task (Jiang et al)

No new data was collected in the direct comparison task here. The data collected by Jiang et al. (2022) being public, it was possible to compare results found in the two previous tasks to direct comparison scores for the exact same set of images. In their study, all images were placed on white backgrounds and resized to a standard size, slightly smaller than the size of the background (600×600 pixels).

### Procedure

#### Visual search task

Observers searched for a target image in a visual environment composed of multiple images. On each trial, the name of the target appeared for 1 second at screen center (for example “elephant”; Figure 1A). Target and distractors were then presented. Participants were to scan the scene until they found the target, to stay fixated on the target once they had found it and to press the space bar. There was no time pressure on finding the target. Each participant performed 180 trials, in a single block that took approximately 45 minutes.

### Triplet odd-one-out task

On each trial, in a separate block, participants were presented with three objects and were asked to click on the odd one out with the mouse pointer. Participants performed 240 trials in a single block that took approximately 30 minutes to complete. The triplet on any given trial was selected pseudo-randomly to distribute the observations evenly across cells in the similarity matrix.

### Direct comparison task (Jiang et al)

Participants were shown two images and had to indicate their similarity on a scale going from 1 (“not at all similar”) to 5 (“highly similar”). There were two similarity conditions: when participants were instructed to use visual similarity to respond; and when they were instructed to use semantic similarity. For comparison with the current study, data from the semantic condition was used (but results did not differ qualitatively when the visual condition was used instead, which is not surprising given that similarity judgments were highly correlated between instructions; Jiang et al. 2022).

## Data analyses

### Similarity matrices

For each of the three tasks, a 60 by 60 similarity matrix was obtained.

For the visual search task, the image count matrix counted the number of times a distractor *j* appeared with a target *i*. The fixation count index counted the number of times a distractor *j* was fixated when target *i* was the search target. Each participant contributed 180 trials to the visual search task, and each trial contained 12 target-distractor pairs; each participant thus contributed data for 2160 pairs. This is less than one trial for each of the 3600 cells. To increase signal to noise ratio, the matrix was summed across the diagonal, assuming that the probability of fixating distractor *j* when searching for target *i* would be similar to the probability of fixating distractor *i* when searching for target *j*. The ratio between the fixation count and image count matrices was the fixation probability matrix. Each cell *(i,j)* contained the probability of fixating a distractor *i*, given target *j* (or conversely). To further increase signal to noise ratio, participant data was pooled before calculating this fixation probability matrix (i.e. a “supersubject”). With this method, each cell of the matrix had an average of 13 unique observations.

For the triplet task, a similar “supersubject” approach was adopted. Each cell *(i,j)* of the similarity matrix contained the probability of selecting neither item *i* nor *j* as the odd-one-out. Each cell had an average of 36 unique observations.

For the direct comparison task, each cell *(i,j)* contained the score provided by participants when that pair was presented. There were at least 10 ratings per cell (pooled across participants).

Matrices were correlated between tasks. To ascertain significance, permutation tests were performed by shuffling item labels for each similarity matrix, and correlating the resulting shuffled matrices. This was done 1000 times, giving rise to a distribution of correlations to which the true correlation could then be compared. All correlational hypotheses were directional: that there would be a positive correlation between similarity as ascertained in all three tasks. Alpha was thus set at α=0.025, and because there were three correlations, Bonferroni-corrected to α=0.008.

### Image-based measures

To determine whether similarity ascertained by scanpaths during visual search, odd-one-out judgments or direct comparisons was driven by image characteristics, two analyses were carried out. Image similarity was estimated in two ways. First, by looking at pixelwise correlations between images. The RGB coordinates for each pixel in an image *i* was correlated with the coordinates for that pixel in image *j* (excluding the white contours). Second, by correlating spatial frequency distributions of images *i* and *j*. The resulting similarity matrices could then be correlated with the behavioral measures obtained in the visual search, triplet and comparison tasks. Because there were 6 correlations, the one-sided alpha level was Bonferroni-corrected to α=0.004.

### Multi-dimensional scaling

The number of dimensions needed to correctly describe a set of points for which only the pairwise distance is known can be estimated by multi-dimensional scaling (Shepard, 1980). Each of the three task similarity matrices was analyzed with MDS. How well the resulting solution (i.e. number of dimensions) preserves the original pairwise distances is given by the stress level, which ranges from 0 to 1. Stress measures the mismatch between the original pairwise distances and the distances in the n-dimensional map; lower stress means the map represents the data more faithfully. Selecting the best number of dimensions that represents the items was done by visual inspection of the scree plot, which relates stress to the number of dimensions. The best number of dimensions is usually selected as the n for which adding further dimensions does not meaningfully reduce stress (the “elbow” method). To quantify whether the three tasks gave similar MDS solutions, and thus that the underlying psychological space was comparable, the Procrustes distance between them was calculated. Procrustes transformation is a set of geometric transformations (translation, rotation, uniform scaling, or a combination) that may change the size, position and/or orientation of a geometric object or set of dots, but not its shape, i.e. the relative locations of vertex points. It is commonly used to quantify the similarity between two shapes, or between two sets of points in a given n-dimensional space. Procrustes distance is the metric that quantifies the smallest distance between the two sets of points after Procrustes transformation. It ranges from 0 (perfect shape correspondence) to 1 (no correspondence).

To determine whether a Procrustes distance estimate was statistically significant, a permutation test was again used. Item labels were randomly shuffled and Procrustes distance was calculated for the shuffled data. In theory the distance between two random sets of points in an n-dimensional space should be 1 (maximal), and it is against this null distribution that observed Procrustes distances can be compared.

### Generative model

A cross-validation approach was adopted in which representational geometry was determined from 80% of the visual search data, following the same analytical steps as described above. The model was then used to predict which item the eyes should fixate first in the remaining 20% of trials. (The analysis omits trials in which the first fixation went to the target.) The true first fixation in the test set was then correlated with the prediction in individual trials. To estimate the variability of that correlation, this was performed 100 times (each time with a different random 80% of the data as the training set). This analysis thus tests the hypothesis that the first saccade goes towards the most similar distractor.

The same analysis was then performed but replacing the most similar distractor by the second most similar distractor, then by the third most similar distractor, and so on. The final analysis tested whether the second saccade was also guided by psychological similarity. The same procedure as described above was performed for the second saccade, to test the hypothesis that the second saccade aimed for the most similar distractor (when it had not been fixated by the first saccade).

Another generative model was based on representational geometry estimated from the two other tasks (triplet odd-one-out and direct comparison). To improve statistical power, normalized similarity matrices from the triplet task and direct comparison were averaged, and this average matrix was used to predict first fixation. There was no need for a cross-validation approach since training and test sets were independent.

For all models, to determine whether the correlations were significant, a null distribution was made by shuffling the labels in the test set, and correlating the resulting shuffled first fixations with the predictions.

## Results

On average, in the visual scanpath task, participants made 8±2 fixations per trial (not counting the initial fixation at screen center). Figure 2A-C shows the similarity matrices for each task.

**Figure 2.**
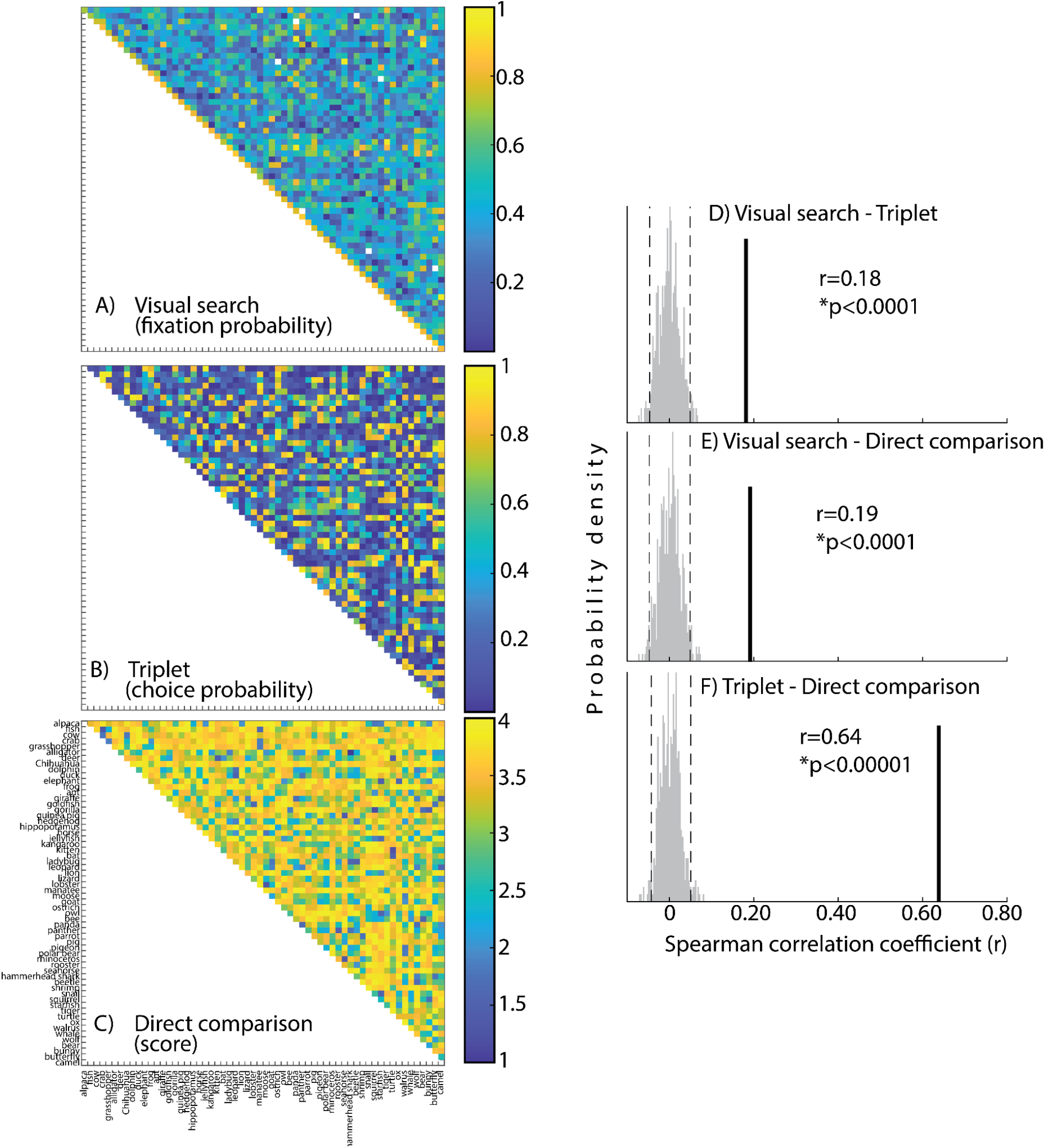
Similarity matrices. The order of the item along the axes is: alpaca, fish, cow, crab, grasshopper, alligator, deer, Chihuahua, dolphin, duck, elephant, frog, ant, giraffe, goldfish, gorilla, guinea pig, hedgehog, hippopotamus, horse, jellyfish, kangaroo, kitten, bat, ladybug, leopard, lion, lizard, lobster, manatee, moose, goat, ostrich, owl, bee, panda, panther, parrot, pig, pigeon, polar bear, rhinoceros, rooster, seahorse, hammerhead shark, beetle, shrimp, snail, squirrel, starfish, tiger, turtle, ox, walrus, whale, wolf, bear, bunny, butterfly, camel. **A)** In the visual search task, similarity is the proportion of times that item *i* (*j)* was fixated when item *j* (*i*) was the target. **B)** In the triplet task, similarity is the proportion of trials in which neither items *i* nor *j* were selected as the odd one out. **C)** The direct comparison score was given by participants on a scale from 0 (not at all similar) to 5 (identical). **D-F)** Correlations between similarity ascertained by the visual search task, the triplet task and direct comparison. Gray distributions are the null distribution obtained by permutation tests, black lines the observed correlations, and insets give the Spearman correlation coefficient and its significance according to the permutation test. Dashed black lines represent the 0.05 significance threshold. D) Correlation between visual search and triplet tasks. E) Between visual search and direct comparison. F) Between the triplet task and direct comparison.

### Correlation between tasks

Similarity across the three tasks correlated (Figure 2D-F). The true correlation between similarity ascertained by scanpaths during visual search and similarity ascertained by the odd one out task was 0.18, which was significant at p<0.001 (Bonferonni-corrected; relative to the null distribution). Visual search behavior and direct comparison scores also correlated (0.19, p<0.001), as did similarity in the odd one out task and direct comparison scores (0.64, p<0.001). Note that correlations with the direct comparison scores, taken from the Jiang et al. dataset, are between-participant; the correlation between visual search and triplet is within-participant.

### Multi-dimensional scaling

Similarity scores from each task were submitted to MDS. To determine the number of dimensions that best describes the data, a scree plot was made relating stress to the number of dimensions. Visual examination of the scree plot in Figure 3A suggests that 2 or 3 dimensions likely suffice. But beyond selecting the appropriate number of dimensions, the real question here was whether the underlying n dimensions were similar between tasks. This was quantified by the Procrustes distance for 3-dimensional MDS (results were qualitatively similar when 2D MDS was considered).

**Figure 3.**
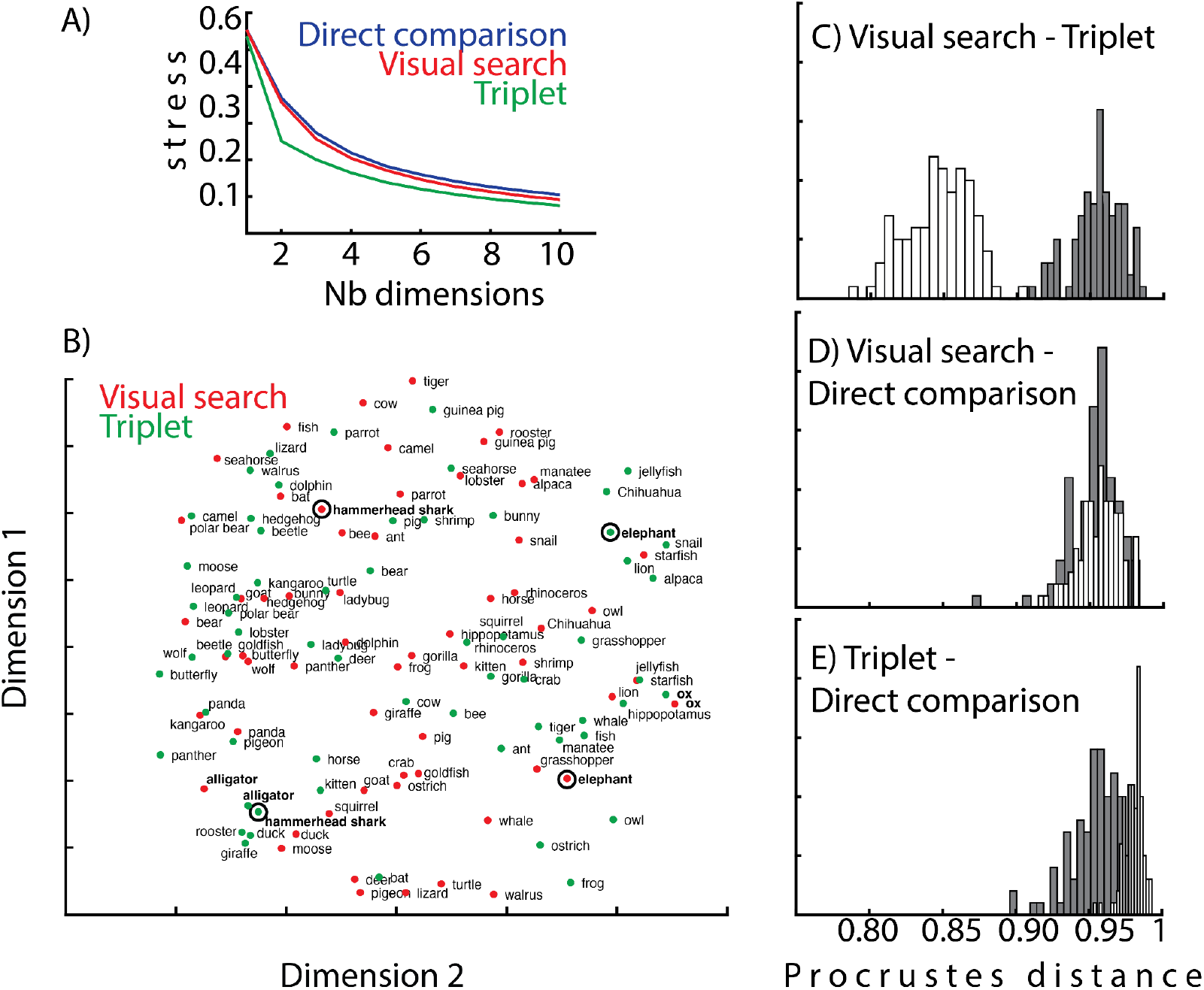
A) Scree plot relating stress to the number of dimensions in the MDS solution, for each of the three tasks. B) Illustration of MDS solutions for the visual search and triplet tasks. Only two of the three dimensions are shown for visualization. C-E) Distribution of observed Procrustes distances and the null distribution for (C) visual search and triplet tasks, (D) visual search and direct comparison tasks and (E) triplet and direct comparison tasks.

Figure 3B shows the MDS solutions for visual search and triplet tasks. Only the first two dimensions are shown for illustrative purposes. For some items, such as “ox” in the bottom lefthand corner, or “alligator” in the righthand corner, the positions are highly similar. For other items, they are not. Compare the hammerhead shark in the visual search task versus in the triplet task (circled in Figure 3B); they have a similar coordinate on dimension 1 but a dissimilar coordinate on dimension 2. The same can be said of elephants. For these two items, it appears that perhaps flipping dimension 2 might align them. This is the principle behind Procrustes transformation Figures 3C-F present the Procrustes distance between visual search, triplet and direct comparison tasks. It is normalized to 1, meaning that the Procrustes distance between visual search and direct comparison, as well as between triplet and comparison, was maximal: the n-dimensional space that best describes each of these tasks was not shared. The Procrustes distance between visual search and triplet tasks was significantly lower than 1, but still rather high (0.85). This may result from the fact that the direct comparison task and the other two tasks were not performed by the same participants, while visual search and triplet were.

### Image-based measures

The similarity between items in the current study could be driven by a number of factors, one of which is visual similarity. Perhaps observers estimate how much each item *looks* like the target and bases their responses on that. Psychological similarity can of course be based on visual characteristics such as color, shape or other such image features, but it likely also includes semantic categories like animacy or functional use. To determine whether the similarity reported here via scanpaths during visual search, odd-one-out judgments or direct comparisons was driven by image characteristics, the behavioral similarity matrices were correlated with the image-based similarity matrices.

Two correlations emerged as significant: between similarity ascertained in the visual search task and SF similarity (r=0.08, Bonferonni-corrected p<0.001 with SF similarity; Figure 4B), and between similarity from direct comparison scores and SF similarity (r=0.08, p=0.012; Figure 4F). These results show that the psychological similarity participants base their behavior on is at least partially driven by low-level visual features. It is important to bear in mind that low-level image similarity does not preclude higher-level, semantic similarity, because exemplars of a common category or nearby categories will, by definition, likely look alike as well.

**Figure 4.**
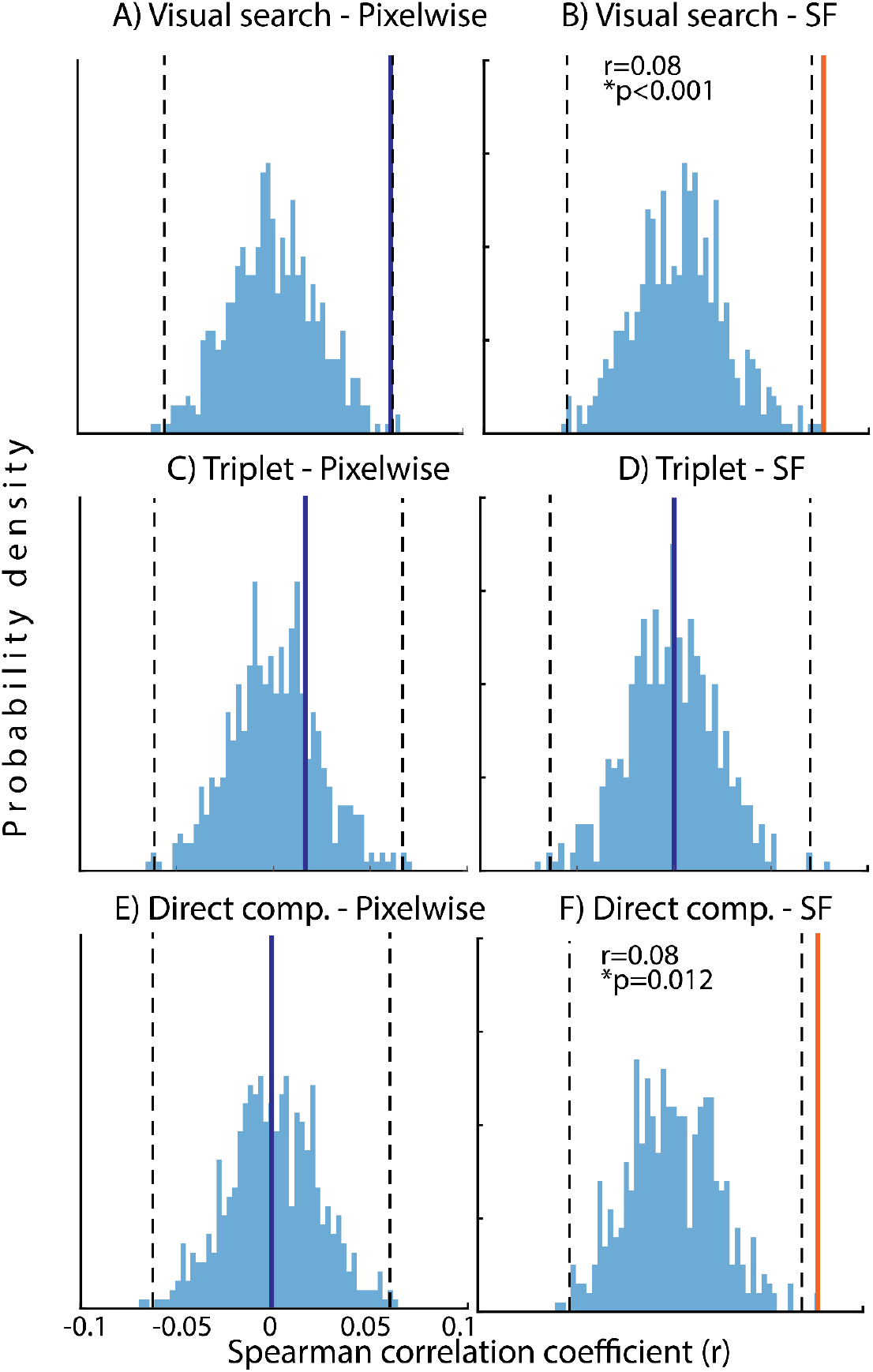
Correlations between similarity ascertained by the visual search task, the triplet task and direct comparison, with image-based similarity measures. Panels A, C and E: pixelwise similarity. Panels B, D and F: spatial frequency distribution similarity. Blue distributions are the null distribution obtained by permutation tests, and insets give the Spearman correlation coefficient and its significance according to the permutation test, when significant. Dashed black lines represent the 0.025 significance threshold. **A)** Correlation between visual search and pixelwise similarity or **(B)** SF similarity. **C)** Correlation between the triplet task and pixelwise similarity or **(D)** SF similarity. **E)** Correlation between direct comparison scores and pixelwise similarity or **(F)** SF similarity.

### Generative model

The psychological similarity based on visual search behavior was estimated on 80% of the data, and tested on the remaining 20%. Figure 5 shows the observed correlation and the null distribution of correlations expected if there was no relation between the most similar distractor and first fixations. The correlation was highly significant: r=0.15 (p<0.0001; red arrow and distribution in Figure 5) meaning that when the first saccade did not go to the target (which actually happened only in a small proportion of trials), it tended to aim for the most similar distractor.

**Figure 5.**
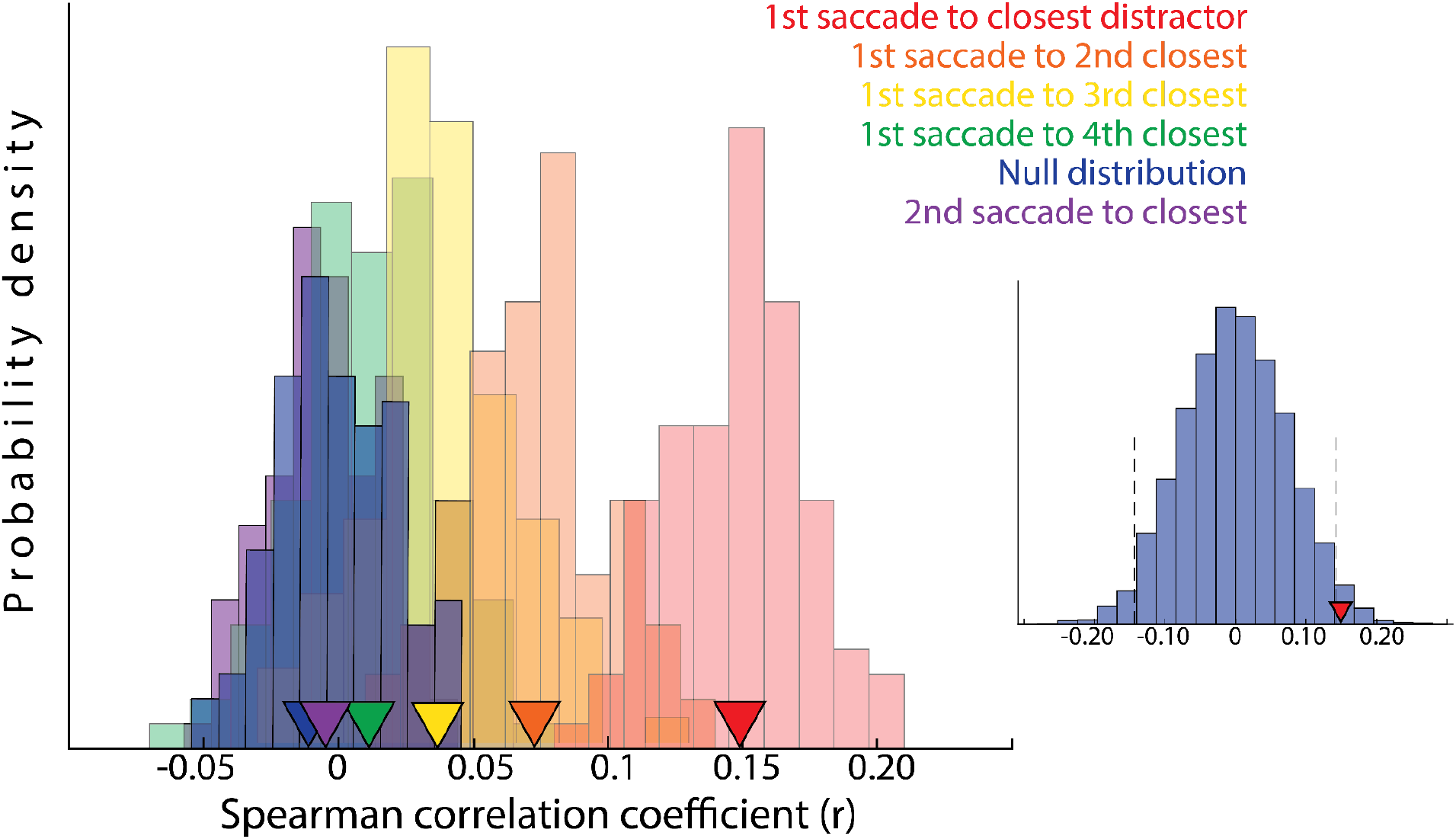
Spearman correlation coefficient between distractors as a function of their similarity to the target (closest, second closest, etc), and the probability of their being fixated first or second. The blue distribution is the null distribution obtained by a permutation test.

The next analysis tested whether there was a graded relationship between the probability of the first saccade aiming for a particular distractor and its similarity to the target. The same procedure as above was applied, but replacing the most similar distractor by the second most similar distractor, then by the third most similar distractor, and so on. The other colored distributions in Figure 5 show these results. There was a significant correlation between predicted and observed first saccades to the second closest distractor (r=0.072, p<0.0001; orange arrow and distribution). The correlation was no longer significant for the third or fourth closest distractors (r=0.037, p=0.08; r=0.011, p=0.28; yellow and green in Figure 5, respectively).

The same procedure was performed for the second saccade, to test the hypothesis that the second saccade aimed for the most similar distractor (when it had not been fixated by the first saccade). There was no evidence in favor of this: the images targeted by second saccades did not correlate with the most similar image in a trial, as can be seen in Figure 5 in which the prediction (in purple) is almost entirely overlapping with the null distribution.

The previous analysis shows that scanpaths during visual search are guided by representational similarity between distractors and targets. This was also true when representational similarity was estimated by the triplet and direct comparison tasks (Figure 5, inset), for first saccades to the closest distractor (r=0.15, p=0.019).

### Functional Visual Field

Average first saccade latency was 177±26 ms (mean±std). This means that the similarity that guides saccades was extracted within less than 200 ms. It is possible that only the items closest to the initial fixation position were estimated. To check this, the next analysis redid the previous analysis (i.e. predicting where first saccades should go if they were driven by representations) but restricting the window in which items were considered. In the previous analysis, similarity between target and distractors was estimated and ranked for all 12 distractors. In the new analysis, similarity was ranked only for targets within a region around screen center (from where the eyes started each trial). The first saccade was thus predicted to aim for the most similar distractor within a window. (This is similar to the notion of the function visual field; see discussion). Figure 6 shows how the correlation evolves with the size of the window. The highest correlations occurred for windows of 2 to 4 dva. The fact that the correlation decreases with larger windows suggests that the role played by distractors outside the window in guiding the eyes is proportional to the efficiency of similarity estimation.

**Figure 6.**
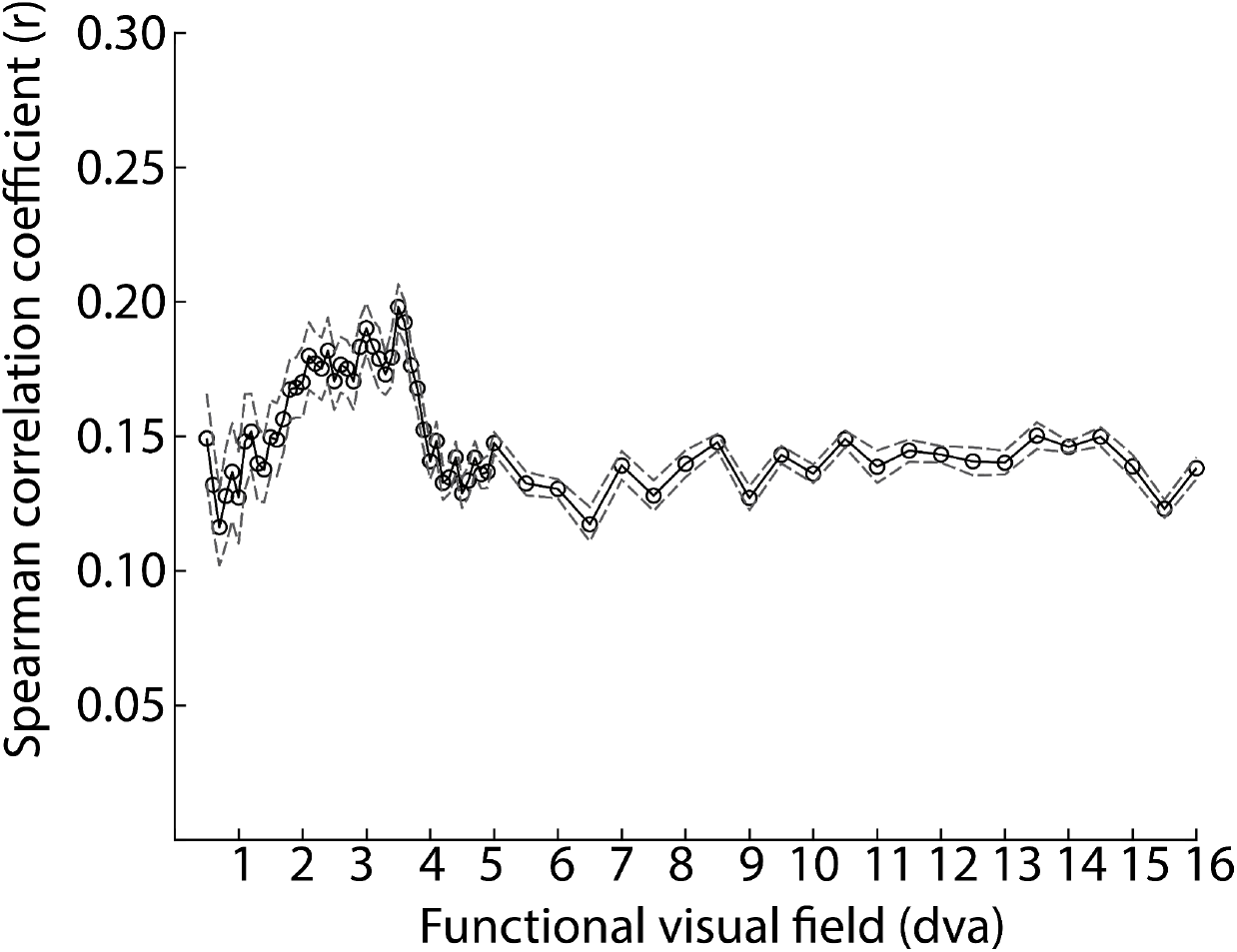
Spearman correlation coefficient between the most similar distractor and the first fixated item, as a function of the size of the window around initial fixation (in dva).

## Discussion

When searching for a visual target amongst distractors, observers scanned the objects according to their similarity with the target. Similarity scores from eye movement patterns are thus an indirect method of inferring the geometry of psychological representations. This indirect method was validated in two ways.

First, convergent validity was obtained by a robust correlation between similarity as ascertained by eye movements in visual search, and similarity ascertained by a triplet odd-one-put task in the same participants. This task has been widely used in the literature to infer representational geometry (e.g. Hebart et al., 2023). Average scanpath similarity also correlated with direct comparison scores obtained in a different set of participants but with the exact same items, published by Jiang et al. (2022). These two measures of convergent validity lend credence to the visual search method. The second validation of the indirect method was criterion validity (the extent to which the method is predictive of future behavior). Representational geometry based on a training subset of visual search, on the triplet task and on direct comparison scores successfully predicted the first item towards which the eyes were drawn in a test subset.

Thus, while representational geometry has traditionally been measured separately from behavior, these results show that it can instead be recovered directly from behavior as it unfolds.

The fact that target-distractor similarity influences visual search performance shows that peripheral information is sufficient to compute values in a priority map or representational space, and that attention and oculomotor guidance must therefore occur before object recognition is complete. In the current study, the similarity of 13 items was estimated in just under 200 ms. Of course, it is possible that only the items closest to the initial fixation position were estimated, and this was examined by restricting the analysis to a particular field of view around the initial fixation. Target-distractor similarity seems to be extracted most efficiently to drive search within a 2-4 dva radius.

Previous research has also shown that eye movements during visual search are guided by an internal representation of the target that is compared to other objects in a search array or in a real-word scene (Wolfe, 2020). Indeed, search becomes slower and less efficient as distractors become more similar in a particular feature to the target (Wolfe, 1994): for example, a red bar among green bars is easy to find, but more difficult when it is among orange bars. Such observations led to the idea that objects are represented in a “priority map” in which candidate objects are compared to a target representation. Visual search and eye movements reveal the evolving match computation.

Visual search can also be guided in real scenes by semantic knowledge about where certain things tend to be (e.g. along a horizontal axis in a street scene if looking for a car; to the top of a tree if looking for a bird, bottom if looking for a rodent; Torralba, Oliva, Castelhano & Henderson, 2006). Henderson & Hayes (2017; 2018) showed that eye movements tended to aim for areas of a scene that had been rated by independent observers as easy to recognize. They called this measure a “meaning map” and showed that when controlling for the correlation between meaning maps and low-level feature maps, only semantic content accounted for unique variance. Hayes & Henderson (2021) further showed that eye movements tended to fixate on areas in scenes that were occupied by objects that were semantically similar to the overall scene meaning, as estimated by a semantic vector-space model based on text corpora.

Although this body of research does not explicitly use the language of representational similarity analysis, the underlying ideas are tightly related. Indeed, this literature usually does not assume that eye movements directly reveal subjective similarity judgments in a pure sense, but rather that fixation probability is better thought of as a similarity-weighted attentional priority signal. If eye movements during visual search are behaviorally observable consequences of similarity computations between internal target templates and candidate scene objects, gaze allocation functions as an indirect measure of representational proximity. A “meaning map” is similar to the idea of representational geometry, but pits semantic and image-based measures against each other. A multi-dimensional representational geometry can have semantic and feature dimensions that jointly determine where the eyes go. Distractors that are similar to the target along a number of possible dimensions are highly activated and are therefore more likely to attract attention and saccades. In all likelihood, which dimensions guide behavior depends on the task (Chan & Hayward, 2014). The task used in the meaning-map experiments was to look at the scenes in preparation for a subsequent memory test (which was not administered). Tasks in studies showing eye movement guidance by visual features often use simple search array with targets defined as features (a red circle, a tilted bar). This task-dependency highlights the importance of developing implicit measures of similarity which can be obtained while participants are doing something else (looking at a scene for a subsequent memory test versus looking for an item in that scene).

If there are multiple maps (semantic, feature), they may need to be combined either in a winner-take-all manner or a weighted average (Liesefeld & Müller, 2019; Buetti et al., 2019). Alternatively, if what guides visual search is a high-dimensional representational signal, it does away with the problem of combining maps.

In sum, the current study shows that eye movements can be used as a read-out of ongoing representational similarity. The recovered similarity space is a latent cognitive representation that gives rise to both eye movements and explicit judgments. That theory doesn’t merely predict that scanpaths correlate with similarity judgments; it specifies why they do. The study thus contributes both the validation of an implicit measure and a proposal of a generative account of visual search behavior. Estimating representational similarity while observers are engaged in another task will pave the way to determining the causal role of representations in behavior.

## Acknowledgments

Gabriela Pincancio & Matteo Lecca, two first-year Master’s student interns in the cognitive science program at Université Paris Cité & Sorbonne Université, collected the data.

## References

Becker, S. I. (2010). Oculomotor capture by colour singletons depends on top-down search goals. Vision Research, 50(2), 211–218. 10.1016/j.visres.2009.11.005

Brainard, D. H. (1997). The Psychophysics Toolbox. Spatial Vision, 10(4), 433–436. 10.1163/156856897X00357

Buetti, S., Cronin, D. A., Madison, A. M., Wang, Z., & Lleras, A. (2019). Four laws of semantic attention: How semantic information affects attention allocation in visual search. Nature Human Behaviour, 3, 408–420.

Chunharas, C., Wolff, M. J., Hettwer, M. D., & Rademaker, R. L. (2024). A gradual transition toward categorical representations along the visual hierarchy during working memory, but not perception. bioRxiv, 2023–05.

Edelman, S. (1998). Representation is representation of similarities. Behavioral and Brain Sciences, 21(4), 449–467.

Henderson, J. M., & Hayes, T. R. (2017). Meaning-based guidance of attention in scenes as revealed by meaning maps. Nature human behaviour, 1(10), 743–747.

Hebart, M. N., Contier, O., Teichmann, L., Rockter, A. H., Zheng, C. Y., Kidder, A., Corriveau, A., Vaziri-Pashkam, M., & Baker, C. I. (2023). THINGS-data: A multimodal collection of large-scale datasets for investigating object representations in human brain and behavior. eLife, 12, e82580. 10.7554/eLife.82580

Henderson, J. M. (2017). Gaze control as prediction. Trends in Cognitive Sciences, 21(1), 15–23.

Henderson, J. M., & Hayes, T. R. (2017). Meaning guides attention in real-world scene images: Evidence from eye movements and meaning maps. Journal of Vision, 17(6), 10.

Jiang, Z., Sanders, D. M. W., & Cowell, R. A. (2022). Visual and semantic similarity norms for a photographic image stimulus set containing recognizable objects, animals and scenes. Behavior research methods, 54(5), 2364–2380.

James, W. (1890). The Principles of Psychology (Vols. 1–2). Henry Holt.

Kleiner, M., Brainard, D. H., Pelli, D. G., Ingling, A., Murray, R., & Broussard, C. (2007). What’s new in Psychtoolbox-3? Perception, 36(ECVP Abstract Supplement), 1–16.

Kriegeskorte, N., Mur, M., & Bandettini, P. A. (2008). Representational similarity analysis-connecting the branches of systems neuroscience. Frontiers in systems neuroscience, 2, 249.

Kriegeskorte, N., & Mur, M. (2012). Inverse MDS: Inferring dissimilarity structure from multiple item arrangements. Frontiers in psychology, 3, 245.

Liesefeld, H. R., & Müller, H. J. (2019). Current directions in visual search. Current Directions in Psychological Science, 28(5), 430–436.

Pelli, D. G. (1997). The VideoToolbox software for visual psychophysics: Transforming numbers into movies. Spatial Vision, 10(4), 437–442.

Rademaker, R. L., Chunharas, C., & Serences, J. T. (2019). Coexisting representations of sensory and mnemonic information in human visual cortex. Nature Neuroscience, 22, 1336–1344.

Shepard, R. N. (1962). The analysis of proximities: Multidimensional scaling with an unknown distance function. Psychometrika, 27, 125–140.

Shepard, R. N. (1980). Multidimensional scaling, tree-fitting, and clustering. Science, 210, 390–398.

Shepard, R. N., & Chipman, S. (1972). Second-order isomorphism of internal representations: Shapes of states. Cognitive Psychology, 3(1), 1–17.

Torralba, A., Oliva, A., Castelhano, M. S., & Henderson, J. M. (2006). Contextual guidance of eye movements and attention in real-world scenes: The role of global features in object search. Psychological Review, 113(4), 766–786.

Walsh, C. R., & Rissman, J. (2023). Behavioral representational similarity analysis reveals how episodic learning is influenced by and reshapes semantic memory. Nature Communications, 14(1), 7548.

Wolfe, J. M. (1994). Guided Search 2.0: A revised model of visual search. Psychonomic Bulletin & Review, 1(2), 202–238.

Wolfe, J. M. (2020). Visual search: How do we find what we are looking for?. Annual review of vision science, 6(1), 539–562.

